# Engineering fluorescent reporters in human pluripotent cells and strategies for live imaging human neurogenesis

**DOI:** 10.1101/2024.05.01.591467

**Authors:** Alwyn Dady, Lindsay Davidson, Nicolas Loyer, Sophie Rappich, Greg Findlay, Timothy Sanders, Jens Januschke, Kate G. Storey

**Author notes:** Division of Molecular, Cell & Developmental Biology, School of Life Sciences, University of Dundee, Dundee DD1 5EH, UK.

## Abstract

Investigation of cell behaviour and cell biological processes underlying human development is facilitated by creation of fluorescent reporters in human pluripotent stem cells, which can be differentiated into cell types of choice. Here we report use of a piggyBac transposon-mediated stable integration strategy to engineer human pluripotent stem cell reporter lines. These express a plasma membrane localised protein tagged with the fluorescent proteins eGFP or mKate2, the photoconvertible nuclear marker H2B-mEos3.2, or the cytoskeletal protein F-tractin tagged with mKate2. Focussing on neural development these lines were used to live image and quantify cell behaviours, including cell cycle progression and cell division orientation in spinal cord rosettes. Further, lipofection-mediated introduction of piggyBac constructs into human neural progenitors labelled single cells and small cell groups within rosettes, allowing individual cell behaviours including neuronal delamination to be monitored. Finally, using the F-tractin-mKate2 hiPSC line, novel actin dynamics were captured during proliferation in cortical neural rosettes. This study presents and validates new tools and techniques with which to interrogate human cell behaviour and cell biology using live imaging approaches.

## Introduction

Live imaging is used to monitor cell behaviour within tissues and to reveal sub-cellular protein dynamics. This includes important cellular processes such as cell cycle progression and changes in cell shape and adhesion as cells differentiate and tissues form. Such observations provide insight into developmental and tissue homeostasis mechanisms and can elucidate disease cell states. The ability to genetically engineer human pluripotent stem cells to express fluorescent proteins facilitates live imaging approaches and allows high resolution investigation of cell behaviour and cell biological processes in human tissue models.

Here we create a series of human pluripotent stem cell reporter lines for live cell imaging using the non-viral plasmid based piggyBac transposon system to introduce fluorescent proteins that localise to a range of subcellular compartments (Ding, et al., 2005, Woodard and Wilson, 2015). This approach involves cloning a sequence of interest into a transposon plasmid so that it is flanked by Inverted Terminal Repeat (ITR) sequences (Figs 1A,B), which allow the piggyBac transposase to then cut and paste the sequence within TTAA-chromosomal sites distributed across the genome (Chen, et al., 2020, Elick, et al., 1996, Lee, et al., 2011, Liu, et al., 2013). Large (Li, et al., 2011) and multiplexed sequences (Kahlig, et al., 2010, Moriarity, et al., 2013) can be introduced, and can also be cleanly excised (Elick, et al., 1996), facilitating cell reprogramming approaches (Woltjen, et al., 2009, Yusa, et al., 2009). The piggyBac transposon system is particularly advantageous for live-cell imaging as it can deliver multiple transgene copies, ranging from 1-10 per haploid genome determined by titration of transposase to transposon ratios (Doherty, et al., 2012). Moreover, different transposases exhibit distinct transduction efficiencies, with hyperactive piggyBac transposase (HyPBase) providing more stable gene expression than native piggyBac or the Tc1/*mariner* transposase *Sleeping Beauty* (Doherty, et al., 2012, Yusa, et al., 2011).

**Figure 1.**
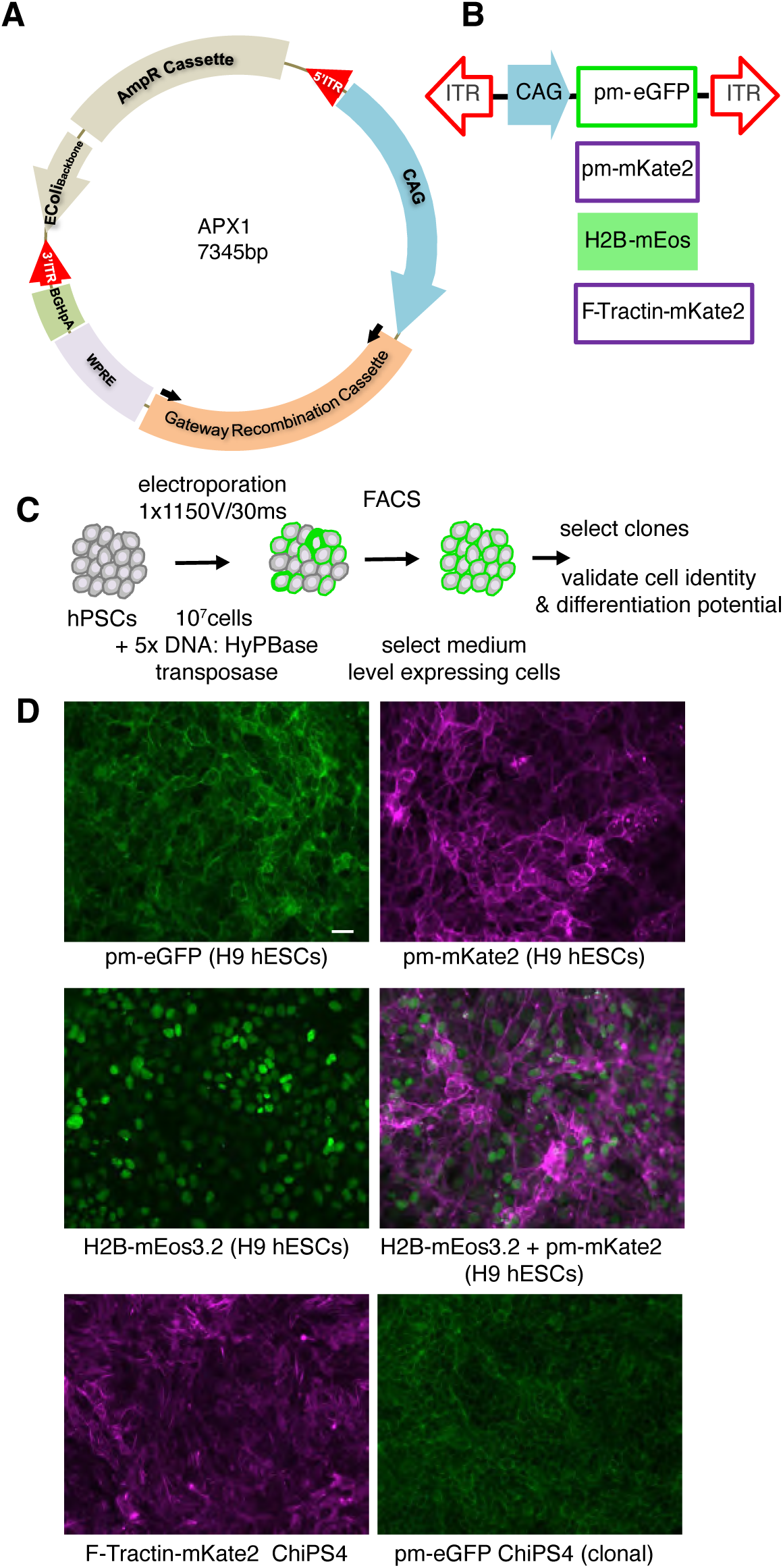
Use of piggyBac to create human pluripotent cell fluorescent reporter lines A) APX1 plasmid used for expression of fluorescent protein fusion constructs highlighting key elements; B) fluorescent-protein reporter constructs for introduction into cells using piggyBac transposase; C) transfection regime and FACS for selection of cells expressing constructs; D) Expression of reporters in human pluripotent cell lines, pm-eGFP, pm-mKate2, H2B-mEos3.2, pm-mKate2+H2B-mEos or F-Tractin-pm-mKate2. Scale bar 50µm.

The robust expression delivered by use of HyPBase may help to ensure good detection sensitivity for fluorescent proteins, in turn reducing imaging exposure times and so potential phototoxicity. This can be further combined with FACS to select moderate expression levels and so minimise potential interference with protein function. Moreover, the transposon system obviates drawbacks of directly engineering endogenous proteins (e.g. using site specific nucleases, TALENS, (Hockemeyer, et al., 2011, Joung and Sander, 2013, Miller, et al., 2011, Wood, et al., 2011) or CRISPR/Cas9, (Charpentier and Doudna, 2013, Doudna and Charpentier, 2014, Jinek, et al., 2012, Pacesa, et al., 2024), which rely on sufficient endogenous promoter strength for fluorescence detection and risk dysfunction of fluorescently tagged endogenous proteins (Roberts, et al., 2017).

Here we use HyPBase to create human pluripotent stem cell reporter lines (or to introduce constructs into cells in *in vitro* generated human tissue), which express fluorescent proteins that localise to the plasma-membrane, chromatin, or actin cytoskeleton. We characterise these lines and transduced tissues and deploy innovative live imaging protocols to capture fundamental characteristics of human neuroepithelial cell behaviour. These tools and approaches allowed analysis of cellular and sub-cellular dynamics during cell cycle progression, including inter-kinetic nuclear migration (IKNM)(Sauer, 1935, Spear and Erickson, 2012) and cell delamination in developing spinal cord, as well as novel F-actin dynamics at the cell membrane and nuclear envelope in cerebral cortex.

## Results and Discussion

### Generation of human pluripotent cell fluorescent reporter lines using the piggyBac-transposon stable integration system

To facilitate analysis of cell behaviour in live tissue derived from human pluripotent stem cells we generated a series of fluorescent reporter hESC or iPSC lines (Dady, et al., 2022) (Figs 1A,B). To monitor cell shape and dynamics we used a plasma membrane (pm) localised protein tagged with eGFP or mKate2 (pm-eGFP or pm-mKate2). To allow monitoring of the cell nucleus we engineered a line expressing H2B-mEos3.2, the latter localising in the nucleus and mEos3.2 having the potential for laser-mediated photo-conversion (Zhang, et al., 2012) and so labelling and tracking of individually selected cells. This would allow clonal analysis in combination with immunofluorescence for cell type specific marker proteins. We also co-transfected H2B-mEos3.2 and pm-mKate2 expressing plasmids to allow monitoring of labelled nuclei and changes in cell shape. Finally, to observe actin cytoskeletal dynamics we selected F-tractin, for its minimal impact on cytoskeletal homeostasis (Belin, et al., 2014), and created an F-tractin-mKate2 cell line.

These reporters were generated using the hESC line H9 (for pm-eGFP, pm-mKate2, H2B-mEos3.2), or hiPSC line ChiPS4 for F-Tractin-mKate2 and a further pm-eGFP line (Dady, et al., 2022) and we deployed the piggyBac transposon-mediated stable integration strategy (Sanders, et al., 2013, Sarkar, et al., 2003, Yusa, et al., 2009, Yusa, et al., 2011). The piggyBac vectors include the fusion protein of interest driven by a CAG promoter. This cargo is then flanked by two Inverted Terminal Repeat (ITR) sequences which allow a hyperactive piggyBac transposase HyPBase to cut and paste cargo within TTAA-chromosomal sites (Figs 1A,B).

The cloning strategy for generating the reporter constructs is described in the Methods and see (Sanders, et al., 2013). Once complete these were introduced with HyPBase to a suspension of hPSCs by electroporation: we determined that transfection of 5 times more piggyBac vector than transposase was critical for efficient stable integration (Figs 1C,D) and see Methods. hPSCs transfected with fusion constructs of interest were then FAC-sorted using their specific fluorescence properties (Fig. S1) to generate polyclonal cell lines selected for expression of easily detectable (medium level) fluorescence for live imaging studies. Single cells could then be selected to make clonal lines that would obviate the potential for cell-to-cell variation to impact cell behaviours captured in live imaging experiments: this was done for pm-eGFP in ChiPS4 (Fig. 1D).

### Using human pluripotent cell fluorescent reporter lines to monitor and quantify cell behaviour in human spinal cord rosettes

These fluorescent reporter hPSC lines allowed the development of *in vitro* assays for monitoring and quantifying human cell behaviour. Here we first focussed on capturing fundamental parameters of human neurogenesis using our established spinal cord rosette assay (Fig. 2A) (Dady, et al., 2022, Verrier, et al., 2018). This involves differentiating hPSCs into Neuro-Mesodermal Progenitor-like cells (NMP-like cells) and subsequently into SOX2-expressing neural progenitors, which self-organise into neural rosettes exhibiting localised expression of apico-basal polarity markers such as NCAD/CDH2 by Day 6 (D6) following NMP-L cell plating (Fig. 2A). This differentiation assay generates neural crest and dorsal neural progenitors, with the first neurons apparent from D8, peak neurogenesis at D14 and gliogenesis onset by ∼ D20 (Dady, et al., 2022). This sequence of events aligns with those documented in human embryonic spinal cord with dorsal neurogenesis at forelimb level commencing by CS12/13 and gliogenesis by CS18 (Dady, et al., 2022, Rayon, et al., 2020).

**Figure 2.**
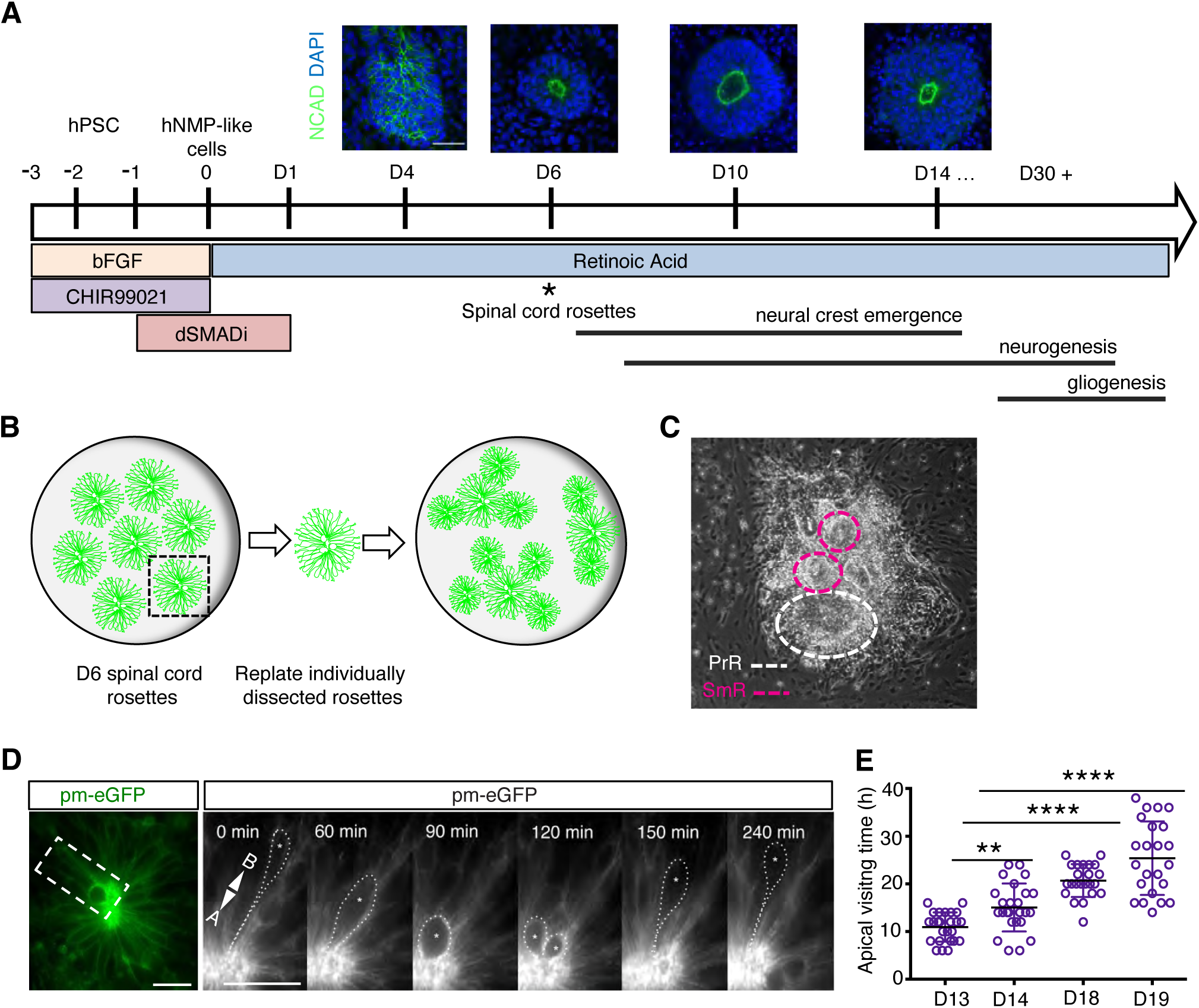
Spinal cord differentiation and rosette generation for live imaging A) Neural differentiation regime after (Verrier, et al., 2018) and neural rosette formation from NMPs (D4-D14), nuclei labelled with DAPI (blue) and apical surface with N-Cadherin (NCAD); B) manual microdissection of D6 primary neural rosettes (PrR), replating and culture to D14/15 and beyond generates; C) small satellite micro-rosettes (SmR) around the PrR, suitable for live imaging; D) Satellite micro-rosette made using pm-eGFP hESC line (Movie 1); E) selected frames from live imaging of SmR following IKNM including mitosis; F) Apical visiting time quantification, revealing increase as development progresses (5 independent experiments for each stage, 25 cells at D13-D14, 23 cells at D18-19), means ± SD, p-values *p<0.05, **p<0.01, *** p<0.001, and ****p<0.0001. Scale bars 50 µm.

We set out to monitor neuroepithelial cell behaviour as cells progressed through the cell cycle during IKNM(Sauer, 1935, Spear and Erickson, 2012). This behaviour is characteristic of pseudo-stratified epithelia and involves movement of the nucleus from the basal to the apical side of a cell in G2 phase, subsequent mitosis at the apical (ventricular) surface and return to the basal surface in G1: here a neuroepithelial cell may then either re-enter S-phase or exit the cell cycle and commence neuronal differentiation. To observe IKNM in spinal cord rosettes we first optimised our assay to address a challenge of longer-term live imaging, increasing thickness of the developing tissue over time (D6 rosettes are typically ∼ 100μm thick). This reduces the ability to maintain cells of interest in focus, even when deploying a long working-distance lens and deconvolution algorithms. To reduce the rosette thickness we developed a re-plating regime in which D6 rosettes were manually micro-dissected and replated on glass-bottom dishes suitable for microscopy (Fig. 2B). This resulted in formation of small satellite micro-rosettes (SmR) surrounding the replated primary rosette (PrR) (Fig. 2C). Satellite micro-rosettes were polarised as indicated by apical actin localisation. Most importantly, SmR were flatter (∼ 50μm thick) allowing longer term high-resolution live cell imaging.

Human spinal cord satellite rosette cells derived from hESCs expressing pm-eGFP or co-expressing pm-mKate and H2B-mEos3.2 were then imaged at 10min intervals for 24h at D13-D14 and D18-D19 (Figs. 2D-E, Movie 1). Apical visiting time (defined as the period during which a basally located nucleus transits to the apical surface, divides and returns to a basal position) was used as an indication of changing cell cycle duration: this period increased from D13-D14 to D18-D19, ∼doubling over this 6-day period (D13, 11h ± 3h; D19, 25h ± 7h (Fig. 2F). This real time measurement confirmed that cell cycle extends as hESC-derived spinal cord progenitors age (Rayon, et al., 2020). These findings further correlate with studies in mouse (Calegari, et al., 2005, Calegari and Huttner, 2003, Takahashi, et al., 1995), and chick (Olivera-Martinez, et al., 2014) embryos showing that neuron-generating divisions, which appear later in development, are characterized by longer IKNM and cell cycle.

### Labelling and live-imaging individual human spinal cord rosette cells using temporally defined lipofection of piggyBac constructs

We further wished to monitor sub-cellular behaviour within the developing neuroepithelium. To achieve this, we devised a strategy to target a mosaic of cells in established neural rosettes using lipofection. PiggyBac constructs and HyPBase transposase were transfected into D8/D9 human spinal cord neural progenitors using lipofectamine (Felgner, et al., 1987)(Fig. 3A). Following further differentiation this approach labelled single cells or small cell groups within rosettes (Figs. 3 B,C).

**Figure 3.**
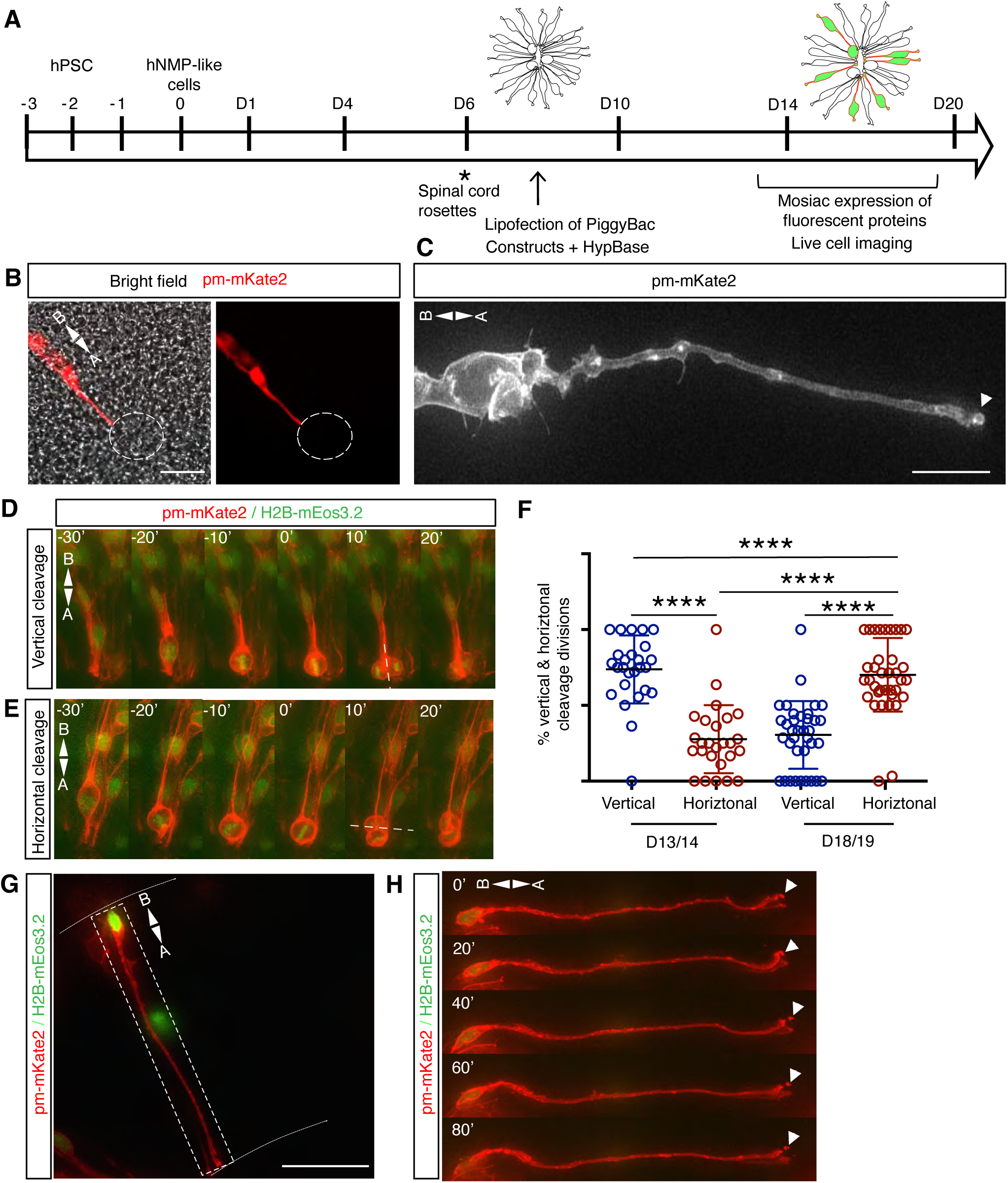
Lipofection-mediated mosaic transfection of spinal cord rosettes A) Regime for lipofection-mediated introduction of piggyBac constructs into spinal cord rosette cells and live-cell imaging; B,C) pm-mKate2 expression revealing individual neuroepithelial cell morphology across apico-basal axis; D,E) Mosaic pm-mKate2+H2B-mEOS3.2 expression allowing live-cell imaging of individual cells and measurement of division symmetry (Movie 2); and F) Percentage of vertical (asymmetric) and horizontal (symmetric) cleavage plane divisions at apical surface in individual rosettes during early and late neurogenesis (5 independent experiments for each stage: D13-14 (181 divisions in 26 rosettes), D18-19 (163 divisions in 36 rosettes) means ± S.E.M; G,H) Live-cell imaging of apical abscission in cell expressing pm-mKate+H2B-mEOS (Movie 3). Scale bars 50 µm.

Lipofection of cells with pm-mKate and H2B-mEos3.2 plasmids revealed cell shape and chromatin configuration and so facilitated analysis of cell division and mitotic spindle orientation. Rosettes containing such cells were cultured from D8/D9 to D13 or D18 and imaged at 10 min intervals for 24-30h. Most divisions between D13 and D14 had vertical cleavage planes, while between D18 and D19 most divisions exhibited horizontal cleavage planes: consistent with a change from neural progenitor cell population expansion to the generation of daughter cells with different, asymmetric, cell fates, such as a neuron and a progenitor (Casas Gimeno and Paridaen, 2022)(Figs 3D-F, Movie 3). These findings demonstrate that division orientation alters as development proceeds in human spinal cord neuroepithelium. This change correlates with the longer apical visiting times observed above and the accumulation of neurons in later stage human spinal cord rosettes (Dady, et al., 2022). To monitor delamination of new-born neurons within rosettes we followed individual cells with a basally located nucleus and a long cellular process inserted into the apical rosette surface. A subset of cells underwent delamination and exhibited abscission of an apical membrane particle in a manner characteristic of the apical abscission mechanism first found in the chicken (Das and Storey, 2014, Kasioulis, et al., 2017)(Figs. 3G, 3H and Movie 5).

Overall, these analyses confirm that expansion of early human spinal cord progenitors is followed by slower cell cycle progression, increased asymmetric cell division and the generation of neurons, and reveal conservation of apical abscission during human neuronal delamination. These findings demonstrate that normal neuroepithelial cell behaviours can be captured using this piggyBac engineering approach to introduce fluorescent reporters.

### Live imaging F-tractin uncovers novel actin dynamics in cortical rosettes

Filamentous actin is a fundamental cytoskeletal component involved in the mechanics of cell division, motility and force transduction within tissues (Davidson and Cadot, 2021, Gunasekaran, et al., 2022, Hyrskyluoto and Vartiainen, 2020), and recent work has identified cell type and cell state specific nuclear roles, e.g. (Okuno, et al., 2020). Within the developing vertebrate neuroepithelium actin mediates IKNM (Norden, et al., 2009, Spear and Erickson, 2012, Yanakieva, et al., 2019) cell adhesion and cell delamination, e.g. (Das and Storey, 2014, Kasioulis, et al., 2017) and intercellular filopodial contacts (Zhang and Scholpp, 2019). Actin dysfunction in human cerebral cortex leads to periventricular heterotopia (Sheen, et al., 2005): it disrupts cell adhesion, proliferation and neuronal delamination and is associated with seizures, dyslexia and psychiatric disorders (Lian and Sheen, 2015). Moreover, the uniquely human fusion gene *CHRFAM7A* acts in part by activating actin in neuronal progenitors, impacting growth cone extension, dendritic spine formation and synaptic clustering (Szigeti, et al., 2023). The mechanisms by which actin regulates such critical processes in the developing human brain have yet to be elucidated.

To analyse actin dynamics, we first established cerebral cortex (cortical) neural rosettes following a dual-SMAD forebrain differentiation protocol after (Chambers, et al., 2009), and rosette structures were generated by D18 (Fig. 4A). Cortical rosettes made with the ChiPS4 parental line and the F-tractin-mKate2 hiPSC line were characterised at D29 for expression of neural progenitor and neuronal markers, and included a few TBR2/EOMES positive cells indicative of forebrain basal progenitors (Figs S2A-D)(detection of TBR2/EOMES was further confirmed in cortical rosettes differentiated with the Wnt signalling antagonist XAV939 Fig.S2E). The F-tractin-mKate2 hiPSC line was then used to investigate actin localisation in fixed tissue and by live-cell imaging using laser-scanning confocal microscopy (D28-29). F-tractin was detected in the neural progenitor cell cortex and was enriched at apical cell junctions and in rounding mitotic cells within rosettes; with similar localisation patterns revealed by phalloidin staining in both the parental ChiPS4 line and F-tractin-mKate2 hiPSC line-derived cortical rosettes (Figs.4B,4C, Movie 4).

**Figure 4.**
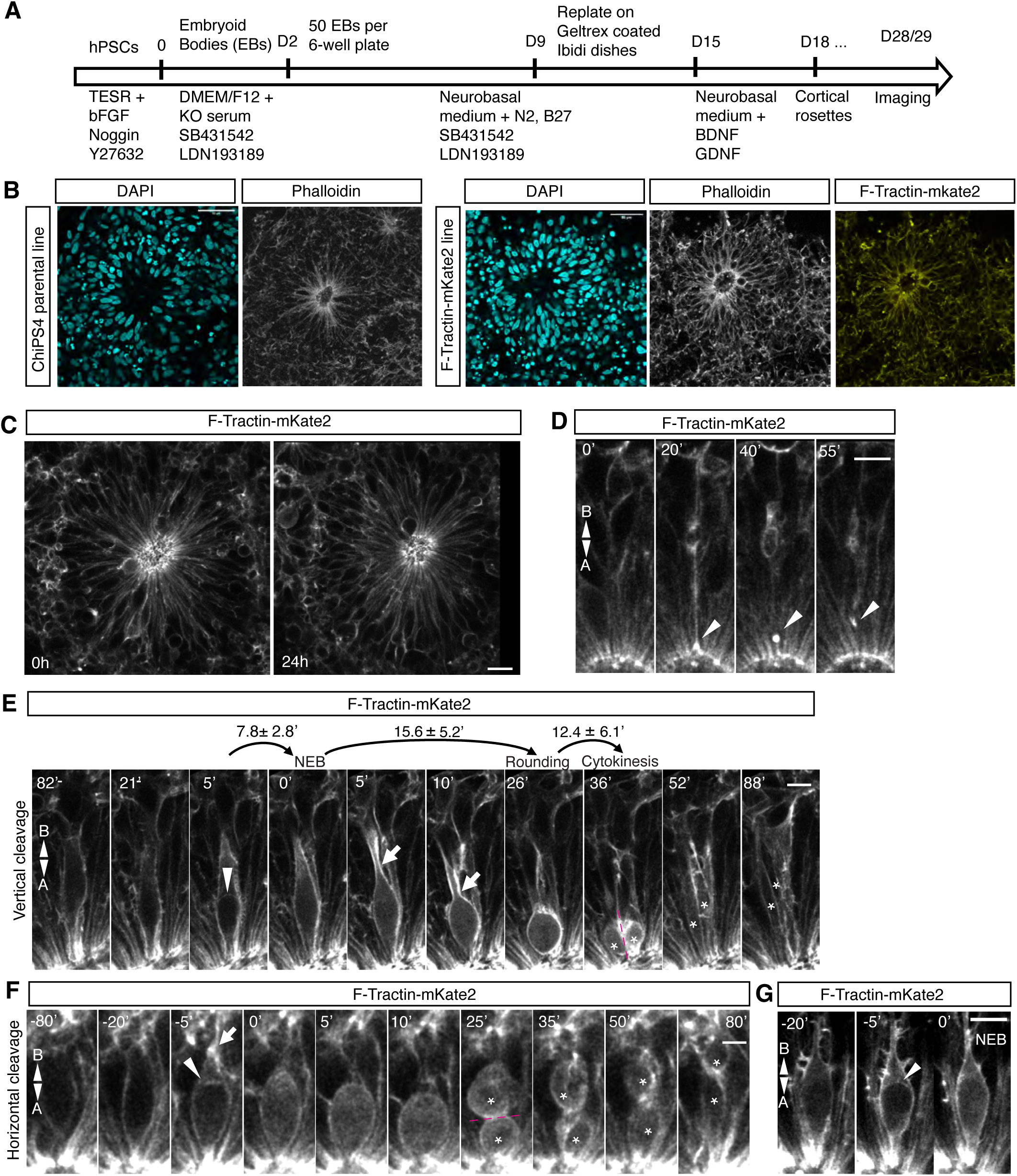
Imaging F-Tractin dynamics in cortical rosettes A) Summary steps of cortical neural rosette differentiation protocol; B) Comparison of F-actin localisation revealed by Phalloidin binding in parental ChiPS4 line (18 rosettes, 1 experiment) and engineered ChiPS4 line expressing F-Tractin-mKate2 (2 independent experiments, 18 rosettes) (phalloidin images maximum-intensity-projections of 4 Z-stack planes at level of lumen); C) Live imaging of cortical rosettes expressing F-Tractin, whole rosette monitored over 24h, note F-Tractin enrichment in apical endfeet in rosette centre (arrowhead) and prominent apical mitoses (Movie 4); D) F-Tractin-mKate2 enrichment at apical tip of a delaminating cell (9 cells in 30 rosettes)(Movie 5); Cleavage plane of divisions assessed in (1 experiment, 113 divisions in 11 rosettes), E) vertical cleavage plane (symmetric) division (103/113 divisions), values above arrows are average time between indicated steps(Movie 6) and F) horizontal cleavage plane (asymmetric) division (10/113 divisions)(Movie 7); G) Neuroepithelial cell in a cortical rosette before and during Nuclear Envelope Breakdown (NEB), representative of observations in 2 independent experiments. A (Apical), B (Basal), Arrowheads: transient F-Tractin enrichment in: D delaminating cell tip, and E,F,G at basal nuclear surface; arrows: F-Tractin-rich basal process; asterisks: daughter cells following cytokinesis. Scale bars: B), 50 µm; C), D) 20 µm, E), F) 10 µm, G) 5 µm.

In movies, apical F-Tractin enrichment was also observed during cell delamination (Fig 4D, Movie 5). Low-level F-Tractin was detected in the cytoplasm but not in the nucleus (Figs 4C-E). Most divisions (97%) took place at the apical surface, with the vast majority (91%) exhibiting a vertical (Fig. 4E, Movie 6) and remaining divisions (9%) a horizontal cleavage plane (Fig. 4F, Movie 7). A small number of basal divisions within the plane of rosettes (Fig.S3, Movie 8) correlated with the few TBR2-expressing cells at this time (Figs.S2A, S2C). Interestingly, during prophase of mitosis, novel F-Tractin was enriched at the basal nuclear membrane prior to Nuclear Envelope Breakdown (NEB) (Figs 4E-G). Actin polymerises around the entire nuclear envelope in mitotic U2OS human cells prior to NEB and associates with nuclear envelope remnants to facilitate chromosome congression and correct segregation (Booth, et al., 2019). This novel actin polymerisation at the basal nuclear surface in cortical progenitors may act similarly, but here act to prevent basal chromosome displacement and so enable apically localised mitosis during IKNM. Following NEB, strong enrichment of F-Tractin was then observed in the basal cell-cortex as the nucleus moved apically and ceased with cell rounding (Figs 4E, 4F). A thin basal process then persisted (Fig 4E, 26’) prior to cytokinesis. Overall, these data reveal established and novel actin localisations and dynamics in human cortical progenitors.

This study demonstrates the use of the HyPBase piggyBac transposase to generate fluorescent protein reporters in human pluripotent cell lines. Efficient stable integration and moderate expression levels were achieved by optimising, i) the quantity and ratio of piggyBac plasmids and transposase and ii) subsequent FACS to exclude high expressing cells, as well as iii) transfection methods, including temporally defined lipofection in hiPSC-derived tissues. The piggyBac system inserts randomly at TTAA sites typically in open chromatin and so is vulnerable to construct silencing during chromatin re-organisation as hiPSCs differentiate. This was mitigated here by use of polyclonal lines and we did not observe silencing in our neural tissue assays, but differentiation for longer periods and into other cell types might lead to loss of expression particularly in clonal lines. Despite this, these approaches will be valuable for cell biological investigation of early differentiation of other cell types from hiPSCs, more sophisticated neural organoids, and explanted tissues. Such assays could be used to investigate environmental stressors (such as toxins or extreme heat) on human cell behaviour. The ability to visualise the actin cytoskeleton and other cell biological components may also facilitate biomechanical studies in human tissues. These reporter cell lines could be further used in pharmacological studies, chemical genetics approaches or functional assays involving acute targeted degradation of proteins using proteolysis targeting chimeras (PROTACs) to elucidate molecular mechanisms underlying fundamental cellular and developmental processes.

## Materials and Methods

### Human pluripotent cell authentication and quality control

Human ESC line H9 was obtained from WiCell and human iPSC line ChiPS4 from Cellartis AB, now Takara Bio Europe. These suppliers provided documentation confirming cell identity. STR DNA profiling for ChiPS4 and ChiPS4-pmGFP (polyclonal), H9 (WA09) confirmed identity of these lines in our laboratory. All cell lines were expanded and banked on arrival. Prior to experiments representative lots of each cell bank were thawed and tested for post-thaw viability and to ensure sterility and absence of mycoplasma contamination. Sterility testing was performed by direct inoculation of conditioned medium into tryptic soya broth and soya bean casein broth and no contamination was observed. Mycoplasma testing was carried out by DAPI staining of fixed cultures and using the mycoalert mycoplasma detection kit (Lonza). We confirm that the cell lines used in this study were free of mycoplasma and aerobic bacteria and fungi.

H9 (WA09) hES cells were supplied at passage 24, the cells were thawed, and cell banks prepared at passage 29: for experiments the cells were used between passage 31 and 41. ChiPS4 hiPS cells were supplied at passage 9 and cell banks prepared at passage 13: for experiments the cells were used between passage 15 and 25. Two passages after thawing cells were checked for uniform expression of pluripotency markers (OCT4, SOX2, NANOG, SSEA-3, SSEA-4 TRA-1-60 and TRA-1-81), absence of differentiation markers (SSEA-1, HNF3-beta, beta-III-tubulin and smooth muscle alpha actinin) by immunofluorescence (details below).

### Human pluripotent cell culture

Human pluripotent stem cells were maintained as feeder-free cultures in TESR medium (Ludwig, et al., 2006) supplemented with bFGF (30ng/mL, Peprotech) and Noggin (10ng/ml, Peprotech) on Geltrex matrix coated plates (10 μg/cm^2^, Life Technologies) at 37°C in a humidified atmosphere of 5% CO_2_ in air. Cells were enzymatically dispersed twice a week using TrypLE select (Life Technologies), counted using a hemacytometer and seeded on Geltrex coated plates at a density of 3e4-5e4 cells/cm^2^. For single-cell passaging, the medium was supplemented by addition of the Rho kinase inhibitor Y-27632 (10 mM, Tocris). All work with hESCs was undertaken in accordance with the UK Stem Cell Bank steering committee guidance (license number SCSC14-29).

### Plasmid expression constructs

The APX1 (Advanced PiggyBac Expression) plasmid was synthetically engineered to harbor improved PiggyBac inverted terminal repeats (ITRs), the cytomegalovirus (CMV) enhancer/chicken Beta-actin (CAG) promoter, woodchuck (hepatitis virus) post regulatory element (WPRE), which increases transgene expression (Higashimoto, et al., 2007), and the Life Technologies Gateway recombination cassette allowing for efficient phiC31-mediated recombination (Sanders, et al., 2013)(Figure 1A). Subcellularly localized fluorescent protein gene cassettes were synthesized by overlap extension PCR and cloned into the Gateway Entry Vector, pENTRDTOPO, and sequence verified. High efficiency transposase-mediated insertion of the PiggyBac transposon cassette was accomplished through transient co-expression of the mammalian optimized PiggyBac transposase (CMV HyPBase, (Yusa, et al., 2011)).

For multicolour labelling of subcellular structures, we utilized monomeric enhanced green fluorescent protein (mEGFP) (gift from Karel Svoboda, Addgene plasmid #18696) (Harvey, et al., 2008) and far-red mKate2 (Evrogen). These proteins were selected for their photostability, low toxicity, high brightness and distinct emission spectra for dual-colour assays making them suitable for live-cell imaging. To label the outer leaflet of the cell membrane, membrane-associated palmitoylated proteins were generated by appending 20-amino acid sequence of ratGAP-43 (MLCCMRRTKQVEKNDEDQKI) to the N-terminus (pm-XFP). To label the cell nucleus, mEos3.2, a green-to-red photoconvertible protein (a gift from Michael Davidson & Tao Xu (Addgene plasmid 54525), was coupled to Histone 2B (H2B) (Zhang, et al., 2012). For labeling of filamentous actin, F-Tractin which contains the inositol triphosphate 3-kinase actin binding domain (Belin, et al., 2014) was coupled to mKate2. Cell lines expressing both H2B-mEos3.2 and pm-mKate2 were then created by co-transfection of each construct and selection for dual expressing cells using FACS.

### hPSCs transfection

H9 hESCs or hIPSCs were transfected using a Neon electroporator using 10 μl tips (Life Technologies). Briefly, cells were dispersed, resuspended in TESR medium and counted as described above. For each transfection 1e6 cells were collected by centrifugation at 300 xg for 2 minutes, washed with 1ml of PBS, then resuspended with 10μl of Neon electroporation buffer R. 1μg of piggyBac vector and 0.2μg of CMV HyPBase vector was then added to the cells in a volume of 2μl. The cells were then electroporated (1150v, 30 mSec, 1 pulse), resuspended in TESR medium supplemented with bFGF, Noggin and Y27632 and plated on Geltrex coated dishes. Cells were maintained as described above then FAC-sorted 10 days after transfection.

### Flow cytometry and cell sorting

Transfected hESCs or hiPSCs were dispersed and re-suspended as described above, then sorted by FACS using a SH800 (Sony) cell sorter as a bulk pool of cells with a narrow gate selected for robustly expressing cells. Quadrant gates used to estimate the percentage of positive cells and level of fluorescence needed were designed based on fluorescence levels detected in the non-transfected control samples. Sorted cells expressing medium levels of fluorescence were expanded and frozen then representative lots of each polyclonal cell line were thawed and tested for mycoplasma contamination using the MycoAlert assay (Lonza), and fungi and aerobic bacteria sterility by inoculating conditioned medium into trypticase soya broth (Sigma Aldrich). No contamination was observed. In some instances (pm-eGFP in ChiPSC4) monoclonal cell lines were made from the polyclonal FACS sorted pools. To make monoclonal cell lines, 400 cells were seeded into 60 mm dishes and cultured until the colonies were 2 – 3 mm in diameter. They were then picked using 3.2 mm cloning discs (Sigma Aldrich) soaked in TrypLE select and seeded into 96 well plates. Individual clones were then expanded in TESR medium and pluripotency, including differentiation in 3 germ layers was confirmed (Supplementary data 1).

### Neural differentiation assays

#### Spinal cord rosette differentiation

This was carried out following (Dady, et al., 2022, Verrier, et al., 2018). Briefly, hESC were plated on Geltrex matrix (20 μg/cm2, Life Technologies) at a density of 4e4 cells/cm^2^ in TSER medium as described and allowed to attach for 24h. Cells were differentiated for 48 hours in Neurobasal medium plus 1X B27, 1xN2, 1x non-essential amino acids and 1x glutamax (all Thermo Fisher) (N2B27 medium) supplemented with 3 µM Chiron99021 (Tocris) and 20 ng/ml bFGF (PeproTech) followed by a further 24 hours in N2B27 medium supplemented with 3 µM Chiron99021, 20 ng/ml bFGF, 10 mM SB431542 (Tocris) and 50 mg/ml Noggin (Peprotech) to obtain NeuroMesodermal progenitors (NMPs). For neural differentiation of NMPs, these cells were then passaged using PBS-EDTA 0.5mM and seeded back at controlled density (1e5 cells/cm2) on Geltrex (20μg/cm2, Life Technologies) in N2B27 medium supplemented with 10 mM SB431542, 50 mg/ml Noggin, 100 nM all-trans retinoic acid (Sigma Aldrich) and 10 mM Y27632 (Tocris). After 24 hours the medium was removed and replaced with N2B27 medium containing 100 nM all-trans retinoic acid for the remainder of the differentiation. As described previously, dorsal spinal cord neural rosettes formed from D6 (Dady, et al., 2022) and medium was refreshed every 3 days.

#### Forebrain/Cortical rosette differentiation

hPSCs were dispersed as described above, counted, and re-suspended in TESR medium supplemented with bFGF (30 ng/ml), Noggin (10 ng/ml) and Y27632 (10 μM) at a concentration of 1e5 cells/ml. 100 μl of cell suspension (1e4 cells) was then added to the required number of wells of 96 well v-bottom plates and the cells pelleted by centrifugation at 300 xg for 5 minutes to form embryoid bodies (EBs). EBs were cultured for 48 hours then collected from the 96 well plates using an 8-channel pipette, pooled and washed twice with DMEM/F12 medium (Gibco). EBs were then resuspended in DMEM/F12 containing 20% knockout serum, 1x glutamax, 1x NEAA, 0.1 mM beta-mercaptoethanol (all Gibco), SB431542 (10 μM, Tocris) and LDN193189 (0.5 μM Sigma Aldrich) and 50 EBs transferred to each well of a low attachment 6-well plate in 2ml of medium to begin differentiation. Every 48 hours 1ml of medium was removed from each well and replaced with Neurobasal (Gibco) containing 1x glutamax, 1x NEAA, 1x N2 (Gibco), 1x B27 (Gibco), SB431542 (10 μM) and LDN193189 (0.5 μM). In some differentiations the Wnt signalling antagonist XAV939 (5 μM) (Tocris) was included from D2 to D6 to increase induction of TBR2/EOMES expressing cells (Figs. S2E,F). On D9 of differentiation the EBs were recovered from the 6 well plates and replated on Geltrex coated (10 μg/cm2) 8 well chamber slides (Ibidi) using 10 – 15 EBs per well in Neurobasal containing 1x glutamax, 1x NEAA, 1x N2, 1x B27, SB431542 (10 μM) and LDN193189 (0.5 μM). The medium was refreshed every 48 hours until D15 of differentiation when it was changed to Neurobasal containing 1x glutamax, 1x NEAA, 1x N2, 1x B27, BDNF (20 ng/ml, Peprotech) and GDNF (20 ng/ml, Peprotech). The medium was then refreshed every 3 days, neural rosettes were evident from D18 of differentiation.

#### Labeling individual cells in human spinal cord rosettes by lipofection

For single cell or small cell group transfection, hESC (H9) cells differentiated into NMP-L cells ( as above) were replated at end of D3 and cultured for 8 to 9 days and then transfected using Lipofectamine 3000 kit (Thermo Fisher) following manufacturer’s instructions with a combination of piggyBac plasmids (pm-eGFP + HypBase or pm-mKate + H2B-mEos3.2 + HypBase) for a final concentration of 2 μg/μl of DNA mix. These lipofected human neural progenitors were then cultured and imaged for the desired period.

#### Immunofluorescence

hPSCs and spinal cord neural rosettes were fixed in 1% paraformaldehyde solution overnight at 4°C. Cortical rosettes were fixed in 4% formaldehyde for 30 min at room temperature. Samples were then washed in PBS and permeabilized with 0.1% Triton X-100 or 0.1% Tween, 0.1% Triton-X in PBS (PBST), blocked with 2% bovine serum albumin (BSA) or 5% donkey serum (as appropriate) for 2h at room temperature. Primary antibodies were diluted as listed below in PBST and samples incubated overnight on a rocker at 4°C. After washing in PBS, appropriate secondary antibodies conjugated to Alexa-fluor 488, 568 or 594 (Life Technologies) were applied for 2h at room temperature. Following further washes in PBS samples were exposed to DAPI (1:1000) for 30 minutes, to visualize nuclei. F-actin cytoskeleton was labeled using Phalloidin coupled with FITC fluorophore (1:1000; ThermoFisher).

Commercially available primary antibodies used to characterize cell types: OCT4 Cell Signaling Technology (C30A3) rabbit mAb (1:400); SOX2 Cell Signaling Technology (D6D9) rabbit mAb (1:400) or R&D Systems (Cat# AF2018, RRID:AB_355110) goat (1:250); NANOG Cell Signaling Technology (D73G4) rabbit mAb (1:400); SSEA-4 Developmental Studies Hybridoma Bank (MC-813-70) mouse mAb (2μg/ml) TRA-1-60 Cell Signaling Technology mouse mAb (1:200);TRA-1-81 Cell Signaling Technology mouse mAb (1:200);SSEA-1 Developmental Studies Hybridoma Bank (MC-480) mouse mAb (2 μg per ml);HNF-3 beta Cell Signaling Technology (D56D6) rabbit mAb (1:400) beta-III-tubulin Sigma Aldrich (SDL.3D10) mouse mAb (1:800); Smooth muscle alpha actinin Developmental Studies Hybridoma Bank (1E12) mouse mAb (2μg/ml); NCad, SIGMA, 1:500, SC-216; TBR2/EOMES 1:200 Abcam 23345, TUJ1, Covance Cat# MMS-435P, mouse monoclonal 1:1000 or Biolegend (Cat# 801202, RRID:AB_2313773) mouse monoclonal 1:1000.TBR1 Abcam (ab31940), RRID:AB_2200219 rabbit pAB (1:200); mKate OriGene TA180091, RRID:AB_2622285 mouse mAB (1:250).

Fluorescence images of fixed samples were taken on an EVOS Cell Imaging System (Thermo Fisher Scientific) using x20 air lense, for hPSCs, on a Leica SP8 confocal for spinal cord rosettes using x20 air lense, while cortical rosettes were imaged using a LEICA SP8 Stellaris confocal microscope (using x40 or 63x NA 1.4 oil immersion objective).

#### Generating satellite human spinal cord rosettes

From NMP cell passaging, transfected cells were cultured in N2B27 containing 100 nM RA (SIGMA) until day 6 when neural rosettes were easily observable. Using a very sharp glass needle (0.29-0.31 mm) (Vitrolife), neural rosettes were individually micro-dissected and gently deposited on a 35 mm glass bottom petri dish (WPI) previously coated with Geltrex (20µg/cm2, Life Technologies). Micro-dissected rosettes were then cultured in N2B27 containing 100nM RA (SIGMA) until the desired stage of observation.

#### Live imaging of human neural rosettes

Cells in satellite rosettes formed from primary micro-dissected spinal cord rosettes or individual lipofected neural progenitors within rosettes were imaged using a DeltaVision Core microscope system, allowing detection of fine cellular detail at low fluorescence levels with minimal exposure time, in a WeatherStation environmental chamber maintained at 37°C. The spinal cord rosette culture medium was buffered with a 5% CO_2_/95% air mix and maintained in a humid chamber. Images were acquired using an Olympus 40x 1.30 NA objective using a Xenon light source and a CoolSnap HQ2 cooled CCD camera (Photometrics). 30-40 optical sections (exposure time, 10-50 milliseconds for each channel, 1024×1024 pixels, 2×2 binning) spaced 1.5μm apart were imaged for each micro-satellite neural rosette at 10-minute intervals for up to 36h. Images were deconvolved and maximum intensity projections of Z-stacks were made using SoftWorx imaging software (Applied Precision) (Das, et al., 2012).

Confocal live imaging of cortical rosettes expressing F-Tractin+mKate2 was undertaken at 37°C with a 5% CO_2_/95% air mix using a Leica SP8 Stellaris confocal microscope or a Zeiss spinning disc confocal microscope with a x63 NA1.4 oil objective. Stacks of 18 slices optical z-section separated by 0.8 µm, covering a 154 × 154 ×14.4 µm region were acquired over multiple positions every 310 seconds for up to 24 hours. We applied a 3D gaussian blur with a sigma of 0.8x 0.8x 0.8 µm.

#### Quantification of cell behaviour

In spinal cord rosettes, apical visiting time was assessed by measuring time taken for a single labelled cell located in the basal region of one spinal cord rosette (rosette periphery) to move to the apical region and divide (rosette centre). To be consistent, observed rosettes were selected by their similar diameter ∼100-150 μm. At least 5 different cells per spinal cord rosette and at least 5 different rosettes were analyzed for each time point (D13, D14, D18, 1D9).

Cell division plane was assessed by measuring the angle of the metaphase plate relative to the apical surface at the lumen wall of the spinal rosette during cell division using Fiji tools. Cleavage plane was considered horizontal when the metaphase plate was perpendicular to the axis of cell division. On the contrary, cleavage plane was considered vertical when the metaphase plate formed an angle of 40-45° to the axis of cell division (with one daughter cell remaining longer at the apical surface). All cells were analysed in 3D. At least 5 different cells per spinal cord rosette and at least 5 different spinal cord rosettes were analyzed for each time point.

In cortical rosettes, we measured the time between cell cycle phases in rosettes expressing F-Tractin-mKate2. Nuclear Envelope Breakdown was identified as the moment when cytoplasmic F-Tractin entered the position held by the nucleus during the previous timepoint. Completion of apical-to-basal migration as the moment when the cell appeared rounded at the apical surface. Cytokinesis onset was defined as the timepoint when the first membrane cortical contraction was detected. Cell division cleavage planes were considered horizontal whenever one daughter cell appeared to have lost all contact with the apical junctions immediately after the completion of cytokinesis. 1 to 21 cells were analysed per rosette amounting to 113 cells in 11 rosettes imaged for 22 hours from D29-D30 were analysed.

#### Statistical analysis

Statistical analysis for spinal cord rosettes was performed using non-parametric Mann-Whitney U test for non-normally distributed data using Prism V8. Results are represented as means ± SEM or SD as indicated in figures and p-values are indicated with *p<0.05, **p<0.01, *** p<0.001, and ****p<0.0001.

## Acknowledgements

The authors thank the University of Dundee Human Pluripotent Stem Cell Facility, Imaging Facility and the Flow Cytometry and Cell Sorting Facility for their contributions to this work.

## Competing interests

The authors declare no competing interests.

## Funding

This work was supported by a Wellcome Trust Investigator award to KGS (WT102817AIA-enhancement) and a Biotechnology and Biological Sciences Research Council grant (BB/V001353/1) to JJ. SR is supported by a Wellcome studentship co-supervised by KS and GF. Initial plasmid generation and cloning experiments were supported by a Children’s Hospital of Pittsburgh of UPMC Research Advisory Committee Grant to AD. The Leica Stellaris confocal microscope was supported by Biotechnology and Biological Sciences Research Council grant BB/T017546/1 and the Leica SP8 by a Wellcome Trust Multi-User Equipment grant (WT101468).

## Data availability

All relevant data can be found within the article and its supplementary information.

## Supplementary information

### Movie legends

#### Live imaging of cell behaviour in human spinal cord rosettes

Movie 1 (Figs 2D,E): nuclear apical-basal visiting in pm-eGFP-expressing spinal cord rosette

Movie 2 (Figs 3D,E): symmetric and asymmetric (final division) in spinal cord rosette following lipofection-mediated mosaic expression of pm-mKate2+H2B-mEos3.2

Movie 3 (Figs 3G,H): Apical delamination of neuroepithelial cell following lipofection-mediated mosaic expression of pm-mKate2+H2B-mEos3.2

#### Live imaging of cell behaviour in human cortical rosettes

Movie 4 (Fig 4A): cell behaviour in F-Tractin-mKate2-expressing cortical rosette

Movie 5 (Fig 4B): Apical delamination of cortical progenitor cell.

Movie 6 (Fig 4C): Apical symmetrical division of cortical progenitor cell. Movie 7 (Fig 4D): Apical asymmetric division of cortical progenitor cell. Movie 8 (Fig 4SD): Basal cortical progenitor cell division. Scale bars: 5 µm Movie 7; 10 µm, Movies 5, 6, 8 (bottom); 20 µm, Movies 4, 8 (top).

### Supplementary Figures

Figure S1 - FASC data for human pluripotent cells expressing fluorescent reporters (related to Figure 1) FASC data for selection of human pluripotent cells expressing fluorescent reporters, showing expression of eGFP in 82.82% of cells, mKate2 in 92.74% of cells, and H2B-mEos3 only in 22.75% of cells, while 13.79% of cells co-expressed pm-Kate and H2B-mEos3.2.

Figure S2 - Characterisation of cortical rosettes (related to Figure 4) Immunofluorescence analysis of cortical rosettes, visualisation of nuclei with DAPI, F-actin with Phalloidin or mKate antibody to detect F-tractin-mKate2, in parental hiPSC line ChiPS4, A) labelling neural (SOX2), and basal (TBR2/EOMES) progenitors, and B) neurons TUJ1; in F-tractin-mKate2 line labelling C) TBR2/EOMES, and D) TUJ1; in parental ChiPS4 line following protocol with XAV939, E) TBR2/EOMES, and SOX2, F) TUJ1. Fifteen rosettes assessed from 1 experiment for each condition (with or without XAV939) in parental ChiPS4 line, 30 rosettes from 2 independent experiments assessed F-tractin-mKate2 line. Scale bars in A-F 50 µm.

Figure S3 - Basal cortical progenitor cell division A) Basal cortical progenitor cell division (within the plane of the rosette, 1 experiment, 3 divisions in 11 rosettes) (movie 8). Scale bars: 20 µm (top) and 10 µm (bottom).

Figure S4 - Pluripotency confirmation of ChiPS4 clonal line pm-eGFP by immunofluorescence Immunofluorescent detection of pluripotency proteins OCT4, NANOG and SOX2 in ChiPS4 clonal line pm-eGFP. Nuclei labelled with DAPI. Scale bar 50µm.

### Supplementary data

Supplementary data 1 - Pluripotency confirmation of ChiPS4 clonal line pm-eGFP by qPCR (Excel file) The clonal ChiPS4 line expressing pm-GFP was quality controlled in 3 technical replicates by qPCR for expression of pluripotency markers *OCT4* and *NANOG* and markers for differentiating endoderm, mesoderm or ectoderm for *FOXA2*, *MSGN1*, *PAX6*. Spontaneous in vitro differentiation of embryoid bodies (in the absence of bFGF) evidenced differentiation into cells representative of the 3 germ layers further confirming pluripotency of this engineered cell line.

## Notes

### Competing Interest Statement

The authors have declared no competing interest.

### Summary of Updates

The Introduction has been revised, including updated references. More detailed documentation of engineered hiPSC lines is now provided in Figures 1 and S4 and in Supplementary data 1. Diagrams and images of cell lines in Figure 1 have been revised for clarity. New data characterising human cortical rosettes is provided in new Figures 4A & 4B and in Figures S2 & S3. Two new authors contributing to human cortical rosette characterisation, Dr Greg Findlay and Ms Sophie Rappich, are now included.

## References

Belin, B. J., Goins, L. M. and Mullins, R. D. (2014). Comparative analysis of tools for live cell imaging of actin network architecture. Bioarchitecture 4, 189–202.

Booth, A. J. R., Yue, Z., Eykelenboom, J. K., Stiff, T., Luxton, G. W. G., Hochegger, H. and Tanaka, T. U. (2019). Contractile acto-myosin network on nuclear envelope remnants positions human chromosomes for mitosis. Elife 8.

Calegari, F., Haubensak, W., Haffner, C. and Huttner, W. B. (2005). Selective lengthening of the cell cycle in the neurogenic subpopulation of neural progenitor cells during mouse brain development. J Neurosci 25, 6533–6538.

Calegari, F. and Huttner, W. B. (2003). An inhibition of cyclin-dependent kinases that lengthens, but does not arrest, neuroepithelial cell cycle induces premature neurogenesis. J Cell Sci 116, 4947–4955.

Casas Gimeno, G. and Paridaen, J. (2022). The Symmetry of Neural Stem Cell and Progenitor Divisions in the Vertebrate Brain. Front Cell Dev Biol 10, 885269.

Chambers, S. M., Fasano, C. A., Papapetrou, E. P., Tomishima, M., Sadelain, M. and Studer, L. (2009). Highly efficient neural conversion of human ES and iPS cells by dual inhibition of SMAD signaling. Nat Biotechnol 27, 275–280.

Charpentier, E. and Doudna, J. A. (2013). Biotechnology: Rewriting a genome. Nature 495, 50–51.

Chen, Q., Luo, W., Veach, R. A., Hickman, A. B., Wilson, M. H. and Dyda, F. (2020). Structural basis of seamless excision and specific targeting by piggyBac transposase. Nat Commun 11, 3446.

Dady, A., Davidson, L., Halley, P. A. and Storey, K. G. (2022). Human spinal cord in vitro differentiation pace is initially maintained in heterologous embryonic environments. Elife 11.

Das, R. M. and Storey, K. G. (2014). Apical abscission alters cell polarity and dismantles the primary cilium during neurogenesis. Science 343, 200–204.

Das, R. M., Wilcock, A. C., Swedlow, J. R. and Storey, K. G. (2012). High-resolution live imaging of cell behavior in the developing neuroepithelium. Journal of visualized experiments: JoVE.

Davidson, P. M. and Cadot, B. (2021). Actin on and around the Nucleus. Trends Cell Biol 31, 211–223.

Ding, S., Wu, X., Li, G., Han, M., Zhuang, Y. and Xu, T. (2005). Efficient transposition of the piggyBac (PB) transposon in mammalian cells and mice. Cell 122, 473–483.

Doherty, J. E., Huye, L. E., Yusa, K., Zhou, L., Craig, N. L. and Wilson, M. H. (2012). Hyperactive piggyBac gene transfer in human cells and in vivo. Hum Gene Ther 23, 311–320.

Doudna, J. A. and Charpentier, E. (2014). Genome editing. The new frontier of genome engineering with CRISPR-Cas9. Science 346, 1258096.

Elick, T. A., Bauser, C. A. and Fraser, M. J. (1996). Excision of the piggyBac transposable element in vitro is a precise event that is enhanced by the expression of its encoded transposase. Genetica 98, 33–41.

Felgner, P. L., Gadek, T. R., Holm, M., Roman, R., Chan, H. W., Wenz, M., Northrop, J. P., Ringold, G. M. and Danielsen, M. (1987). Lipofection: a highly efficient, lipid-mediated DNA-transfection procedure. Proc Natl Acad Sci U S A 84, 7413–7417.

Gunasekaran, S., Miyagawa, Y. and Miyamoto, K. (2022). Actin nucleoskeleton in embryonic development and cellular differentiation. Curr Opin Cell Biol 76, 102100.

Harvey, C. D., Yasuda, R., Zhong, H. and Svoboda, K. (2008). The spread of Ras activity triggered by activation of a single dendritic spine. Science 321, 136–140.

Higashimoto, T., Urbinati, F., Perumbeti, A., Jiang, G., Zarzuela, A., Chang, L. J., Kohn, D. B. and Malik, P. (2007). The woodchuck hepatitis virus post-transcriptional regulatory element reduces readthrough transcription from retroviral vectors. Gene Ther 14, 1298–1304.

Hockemeyer, D., Wang, H., Kiani, S., Lai, C. S., Gao, Q., Cassady, J. P., Cost, G. J., Zhang, L., Santiago, Y., Miller, J. C., et al. (2011). Genetic engineering of human pluripotent cells using TALE nucleases. Nat Biotechnol 29, 731–734.

Hyrskyluoto, A. and Vartiainen, M. K. (2020). Regulation of nuclear actin dynamics in development and disease. Curr Opin Cell Biol 64, 18–24.

Jinek, M., Chylinski, K., Fonfara, I., Hauer, M., Doudna, J. A. and Charpentier, E. (2012). A programmable dual-RNA-guided DNA endonuclease in adaptive bacterial immunity. Science 337, 816–821.

Joung, J. K. and Sander, J. D. (2013). TALENs: a widely applicable technology for targeted genome editing. Nat Rev Mol Cell Biol 14, 49–55.

Kahlig, K. M., Saridey, S. K., Kaja, A., Daniels, M. A., George, A. L., Jr. and Wilson, M. H. (2010). Multiplexed transposon-mediated stable gene transfer in human cells. Proc Natl Acad Sci U S A 107, 1343–1348.

Kasioulis, I., Das, R. M. and Storey, K. G. (2017). Inter-dependent apical microtubule and actin dynamics orchestrate centrosome retention and neuronal delamination. Elife 6.

Lee, C. Y., Li, J. F., Liou, J. S., Charng, Y. C., Huang, Y. W. and Lee, H. J. (2011). A gene delivery system for human cells mediated by both a cell-penetrating peptide and a piggyBac transposase. Biomaterials 32, 6264–6276.

Li, M. A., Turner, D. J., Ning, Z., Yusa, K., Liang, Q., Eckert, S., Rad, L., Fitzgerald, T. W., Craig, N. L. and Bradley, A. (2011). Mobilization of giant piggyBac transposons in the mouse genome. Nucleic Acids Res 39, e148.

Lian, G. and Sheen, V. L. (2015). Cytoskeletal proteins in cortical development and disease: actin associated proteins in periventricular heterotopia. Front Cell Neurosci 9, 99.

Liu, X., Li, N., Hu, X., Zhang, R., Li, Q., Cao, D., Liu, T., Zhang, Y. and Liu, X. (2013). Efficient production of transgenic chickens based on piggyBac. Transgenic Res 22, 417–423.

Ludwig, T. E., Levenstein, M. E., Jones, J. M., Berggren, W. T., Mitchen, E. R., Frane, J. L., Crandall, L. J., Daigh, C. A., Conard, K. R., Piekarczyk, M. S., et al. (2006). Derivation of human embryonic stem cells in defined conditions. Nat Biotechnol 24, 185–187.

Miller, J. C., Tan, S., Qiao, G., Barlow, K. A., Wang, J., Xia, D. F., Meng, X., Paschon, D. E., Leung, E., Hinkley, S. J., et al. (2011). A TALE nuclease architecture for efficient genome editing. Nat Biotechnol 29, 143–148.

Moriarity, B. S., Rahrmann, E. P., Keng, V. W., Manlove, L. S., Beckmann, D. A., Wolf, N. K., Khurshid, T., Bell, J. B. and Largaespada, D. A. (2013). Modular assembly of transposon integratable multigene vectors using RecWay assembly. Nucleic Acids Res 41, e92.

Norden, C., Young, S., Link, B. A. and Harris, W. A. (2009). Actomyosin is the main driver of interkinetic nuclear migration in the retina. Cell 138, 1195–1208.

Okuno, T., Li, W. Y., Hatano, Y., Takasu, A., Sakamoto, Y., Yamamoto, M., Ikeda, Z., Shindo, T., Plessner, M., Morita, K., et al. (2020). Zygotic Nuclear F-Actin Safeguards Embryonic Development. Cell Rep 31, 107824.

Olivera-Martinez, I., Schurch, N., Li, R. A., Song, J., Halley, P. A., Das, R. M., Burt, D. W., Barton, G. J. and Storey, K. G. (2014). Major transcriptome re-organisation and abrupt changes in signalling, cell cycle and chromatin regulation at neural differentiation in vivo. Development 141, 3266–3276.

Pacesa, M., Pelea, O. and Jinek, M. (2024). Past, present, and future of CRISPR genome editing technologies. Cell 187, 1076–1100.

Rayon, T., Stamataki, D., Perez-Carrasco, R., Garcia-Perez, L., Barrington, C., Melchionda, M., Exelby, K., Lazaro, J., Tybulewicz, V. L. J., Fisher, E. M. C., et al. (2020). Species-specific pace of development is associated with differences in protein stability. Science 369.

Sanders, T. A., Llagostera, E. and Barna, M. (2013). Specialized filopodia direct long-range transport of SHH during vertebrate tissue patterning. Nature 497, 628–632.

Sarkar, A., Sim, C., Hong, Y. S., Hogan, J. R., Fraser, M. J., Robertson, H. M. and Collins, F. H. (2003). Molecular evolutionary analysis of the widespread piggyBac transposon family and related “domesticated” sequences. Mol Genet Genomics 270, 173–180.

Sauer, F. C. (1935). Mitosis in the neural tube. J. Comp. Neurol. 62, 377–405.

Sheen, V. L., Jansen, A., Chen, M. H., Parrini, E., Morgan, T., Ravenscroft, R., Ganesh, V., Underwood, T., Wiley, J., Leventer, R., et al. (2005). Filamin A mutations cause periventricular heterotopia with Ehlers-Danlos syndrome. Neurology 64, 254–262.

Spear, P. C. and Erickson, C. A. (2012). Interkinetic nuclear migration: a mysterious process in search of a function. Dev Growth Differ 54, 306–316.

Szigeti, K., Ihnatovych, I., Rosas, N., Dorn, R. P., Notari, E., Cortes Gomez, E., He, M., Maly, I., Prasad, S., Nimmer, E., et al. (2023). Neuronal actin cytoskeleton gain of function in the human brain. EBioMedicine 95, 104725.

Takahashi, T., Nowakowski, R. S. and Caviness, V. S., Jr. (1995). The cell cycle of the pseudostratified ventricular epithelium of the embryonic murine cerebral wall. J Neurosci 15, 6046–6057.

Verrier, L., Davidson, L., Gierlinski, M., Dady, A. and Storey, K. G. (2018). Neural differentiation, selection and transcriptomic profiling of human neuromesodermal progenitor-like cells in vitro. Development 145.

Woltjen, K., Michael, I. P., Mohseni, P., Desai, R., Mileikovsky, M., Hamalainen, R., Cowling, R., Wang, W., Liu, P., Gertsenstein, M., et al. (2009). piggyBac transposition reprograms fibroblasts to induced pluripotent stem cells. Nature 458, 766–770.

Wood, A. J., Lo, T. W., Zeitler, B., Pickle, C. S., Ralston, E. J., Lee, A. H., Amora, R., Miller, J. C., Leung, E., Meng, X., et al. (2011). Targeted genome editing across species using ZFNs and TALENs. Science 333, 307.

Woodard, L. E. and Wilson, M. H. (2015). piggyBac-ing models and new therapeutic strategies. Trends Biotechnol 33, 525–533.

Yanakieva, I., Erzberger, A., Matejcic, M., Modes, C. D. and Norden, C. (2019). Cell and tissue morphology determine actin-dependent nuclear migration mechanisms in neuroepithelia. J Cell Biol 218, 3272–3289.

Yusa, K., Rad, R., Takeda, J. and Bradley, A. (2009). Generation of transgene-free induced pluripotent mouse stem cells by the piggyBac transposon. Nat Methods 6, 363–369.

Yusa, K., Zhou, L., Li, M. A., Bradley, A. and Craig, N. L. (2011). A hyperactive piggyBac transposase for mammalian applications. Proc Natl Acad Sci U S A 108, 1531–1536.

Zhang, C. and Scholpp, S. (2019). Cytonemes in development. Curr Opin Genet Dev 57, 25–30.

Zhang, M., Chang, H., Zhang, Y., Yu, J., Wu, L., Ji, W., Chen, J., Liu, B., Lu, J., Liu, Y., et al. (2012). Rational design of true monomeric and bright photoactivatable fluorescent proteins. Nat Methods 9, 727–729.

