## Supplementary figures and images for "Engineering fluorescent reporters in human pluripotent cells and strategies for live imaging human neurogenesis"

### Figure S1

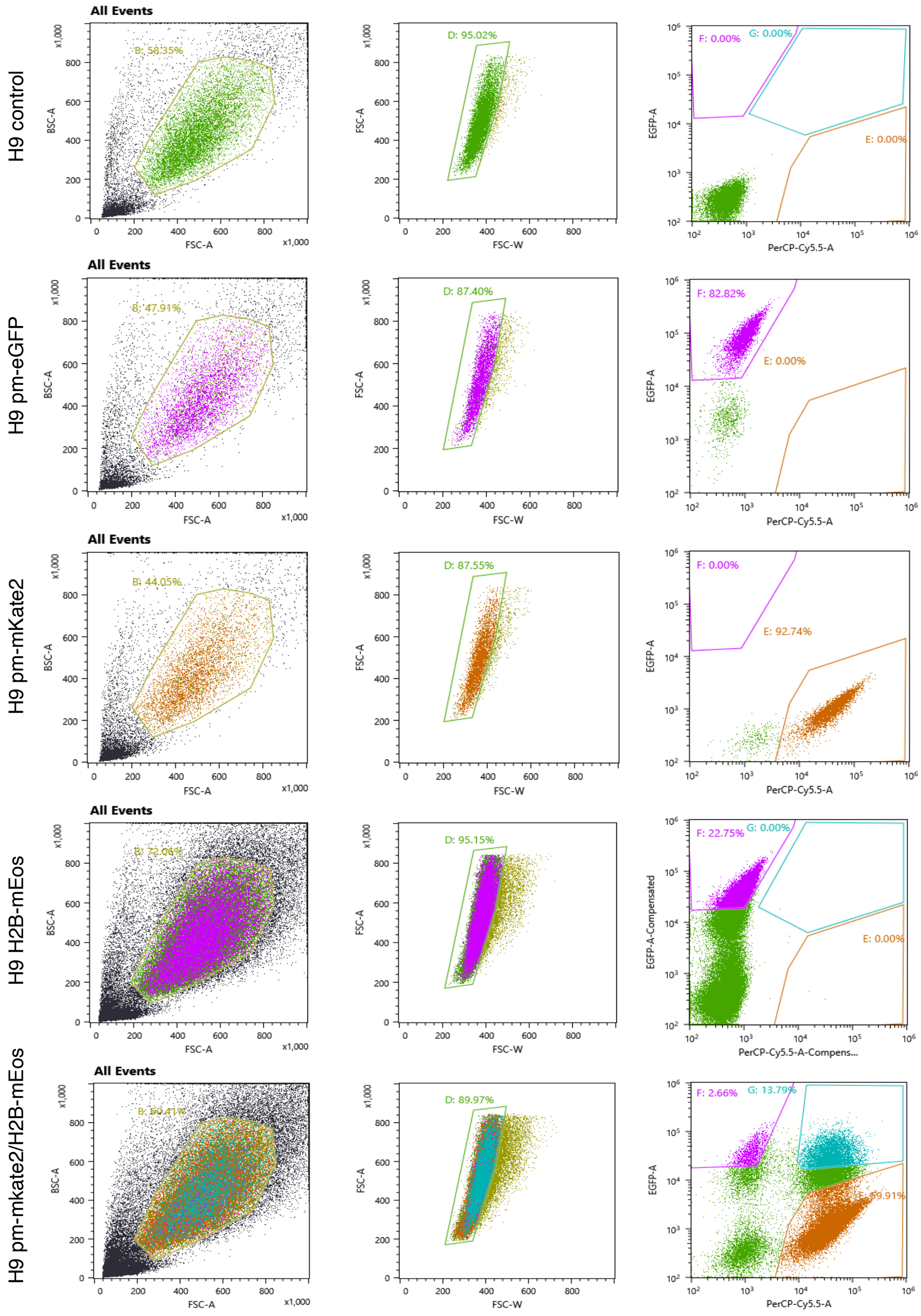


Figure S1 Dady et al

### Figure S2

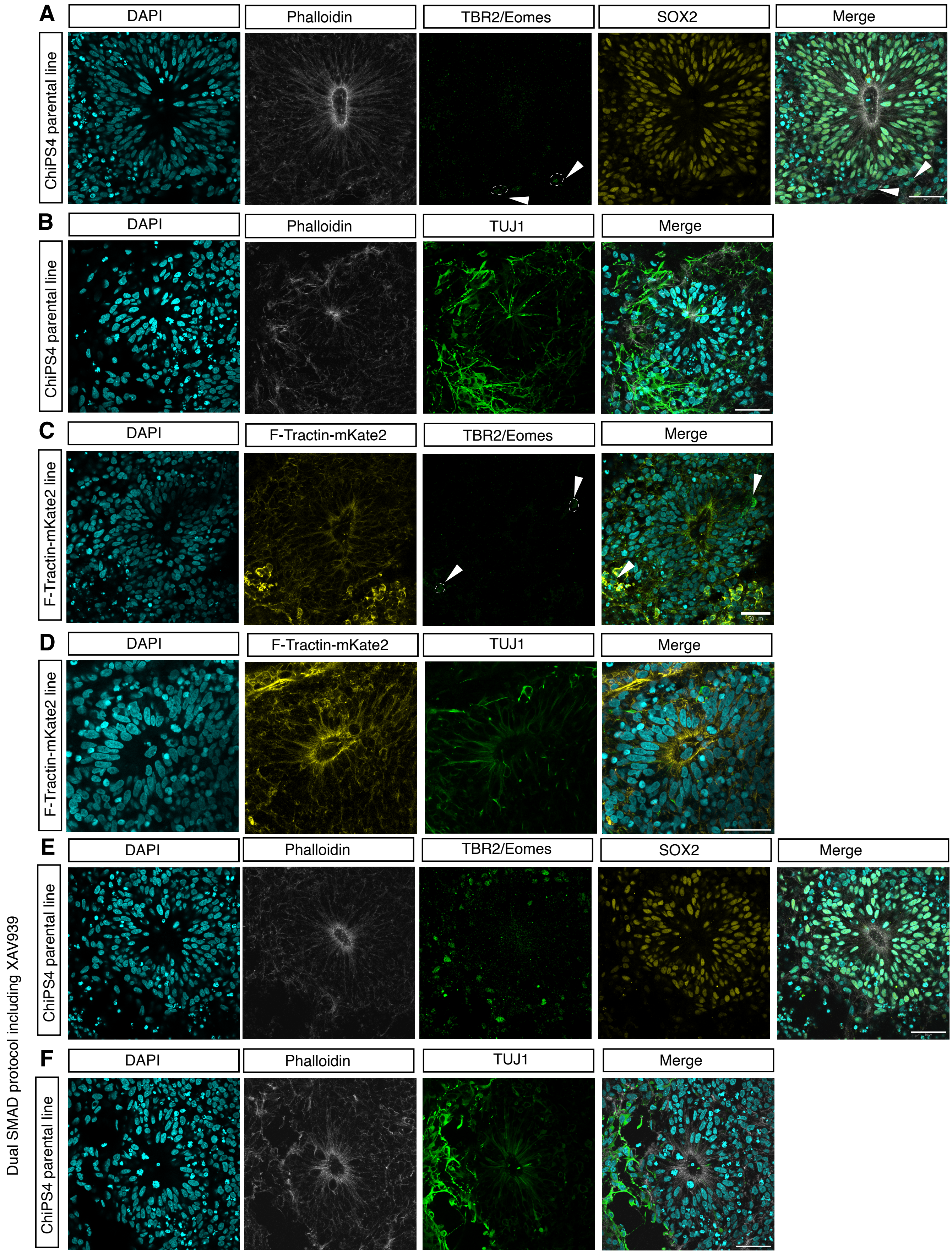


Figure S2 Dady et al

### Figure S3

**A**

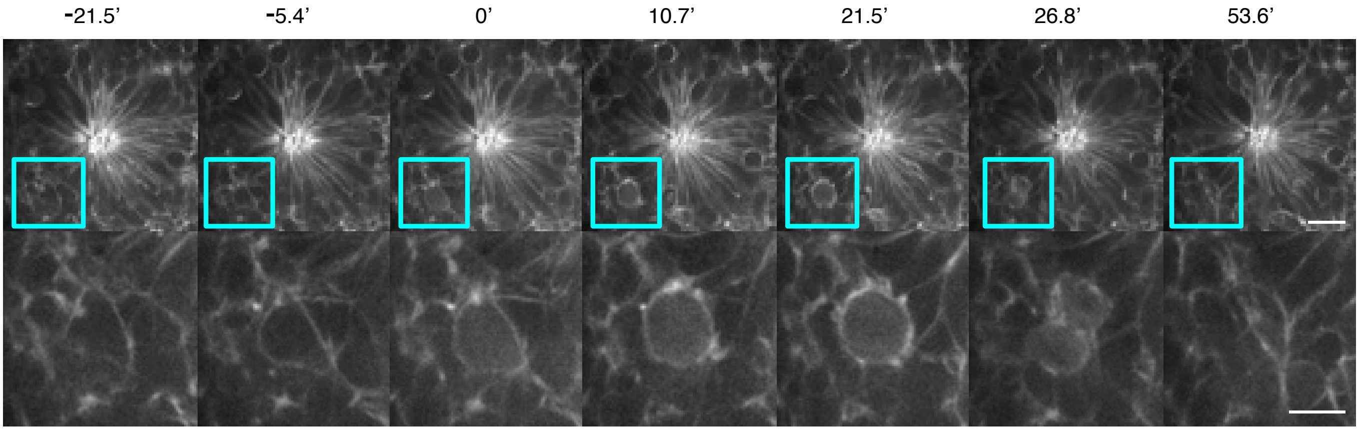


Figure S3 Dady et al

### Figure S4

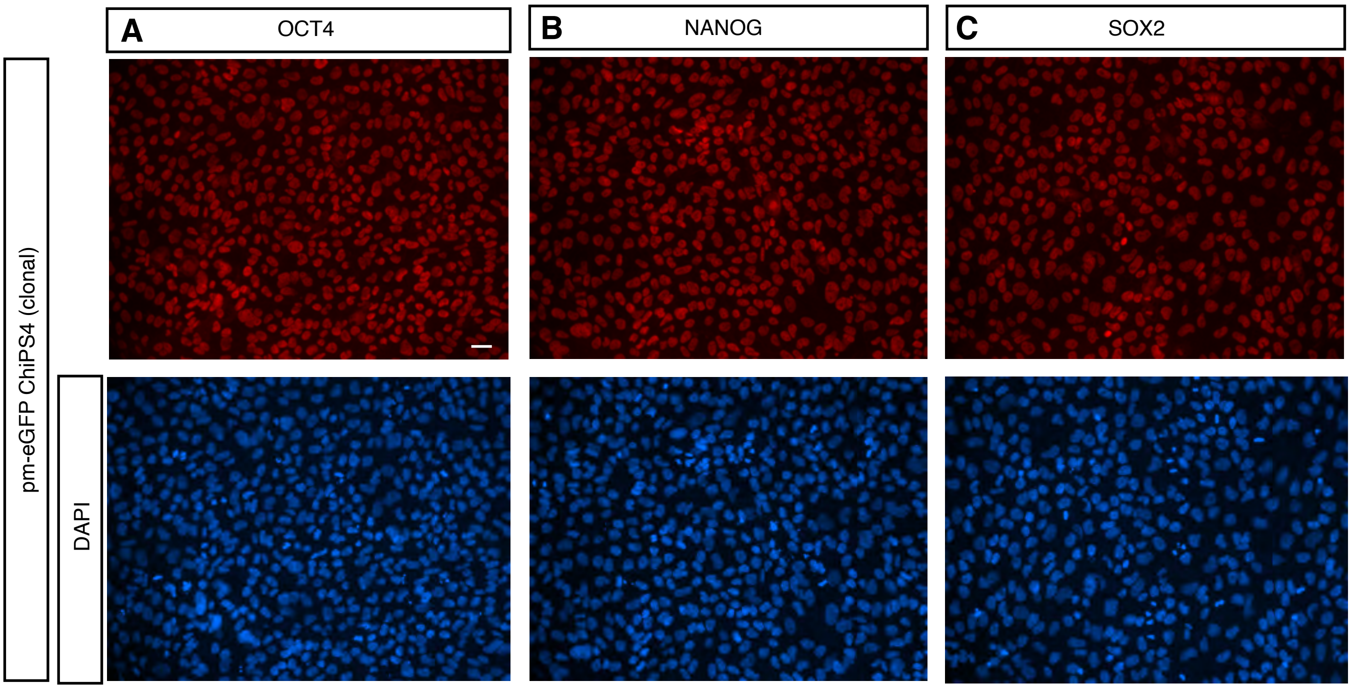


Figure S4 Dady et al
